# Golgi compaction facilitates microtubule nucleation to drive adult vertebrate peripheral neuron regeneration

**DOI:** 10.1101/2024.08.12.607597

**Authors:** Alice E Mortimer, Adam J Reid, Raman M Das

## Abstract

Peripheral neurons regenerate in response to injury, but their cell intrinsic processes are poorly understood and seldom sufficient to effect clinical restoration. Using a novel assay for high-resolution live imaging of regenerating adult vertebrate neurons, we identify acute fragmentation and rapid recompaction of the somatic Golgi as a key driver of peripheral neuron regeneration, implicating this organelle as a therapeutic target in an area of clinical unmet need. Compaction of the fragmented Golgi facilitates stepwise recruitment of the microtubule nucleation factors AKAP9 and γ-tubulin within a discrete period that corresponds with acentrosomal Golgi-mediated microtubule nucleation. Furthermore, disruption of AKAP9 or γ-tubulin recruitment compromises microtubule nucleation leading to impaired regeneration. Crucially, these mechanisms are conserved in the contexts of *in vivo* rat sciatic nerve transection and in primary human peripheral neurons. This work transforms our understanding of the cell intrinsic mechanisms that render injured peripheral neurons competent to initiate axon regeneration.

## Introduction

Regeneration following peripheral nerve injury (PNI) requires extension of axons upwards of one metre to reach the target tissue (1). Current clinical interventions are restricted to surgical repair at the site of injury to provide a permissive environment for regeneration; however, reliance upon innate regenerative capacity is never sufficient for meaningful restoration of function in major nerves (1,2). Indeed, the fundamental cellular mechanisms driving peripheral neuron regeneration remain unknown, impeding the development of new strategies to accelerate this process.

Axon extension is dependent on rearrangement of the neuronal cytoskeleton, of which microtubules are an essential constituent playing vital roles in cellular structure, determining cell polarity, and allowing for bidirectional cargo transport (3). Following PNI, nascent microtubule nucleation presumably drives axon regeneration, yet mature neurons lack a functional centrosome, which acts as the main microtubule organising centre (MTOC) in most cell types; therefore, an alternative cellular compartment must be responsible for microtubule nucleation in regenerating adult neurons (4). Several distinct subcellular compartments have been demonstrated to possess MTOC capacity, including the plasma membrane, the nuclear membrane and the Golgi apparatus (5-8). Importantly, the Golgi apparatus has been implicated as a platform for microtubule nucleation in several cellular contexts (9-13) and microtubule nucleation from Golgi outposts shape cells with unusual morphologies, including neurons and oligodendrocytes (10). In addition, the somatic Golgi has a well-described interplay with microtubules as a means of fulfilling vital roles in polarised cell transport and migration (14,15). The Golgi is therefore a compelling candidate to act as the MTOC in the context of regeneration following PNI.

Here, we demonstrate a conserved neuronal mechanism driving initiation of regeneration in response to PNI across rat and human species, *in vitro* and *in vivo*. Using adult dorsal root ganglia (DRG) sensory neuron injury models which addresses a major unmet clinical challenge to restore sensory function, we observe somatic Golgi fragmentation followed by rapid compaction, before stepwise recruitment of the key microtubule nucleating factors AKAP9 followed by γ-tubulin. Disruption of Golgi compaction prevents recruitment of AKAP9 and γ-tubulin, leading to a reduction in microtubule nucleation and failure of axon regeneration. Furthermore, disruption of AKAP9 recruitment to the Golgi results in reduced γ-tubulin recruitment and ultimately compromises the innate ability to regenerate axons following neuronal injury. These results together demonstrate that compaction of the Golgi and AKAP9 mediated γ-tubulin recruitment to the compacted Golgi drives microtubule nucleation following PNI to facilitate initiation of axon regeneration.

## Results

### Peripheral neuron injury induces Golgi fragmentation and subsequent compaction leading to emergence of microtubules from the Golgi

To investigate the cellular dynamics of axon regeneration following PNI, we induced acute axotomy *in vitro* by dissecting adult rat DRGs from the animal, followed by enzymatic and mechanical dissociation to obtain single neurons. Cells were then cultured and transfected with a plasmid expressing GFP-GPI to label cell membranes and subjected to live imaging using high-resolution widefield timelapse microscopy from 24 hours post-injury. This revealed onset of axon regeneration at 24-30 hours post-injury (21/31 cells, 8 animals). During time-lapse imaging up to 40 hours post-injury, 14/31 cells established a dominant axonal extension, and 7/31 cells established the typical pseudo-unipolar morphology of DRG neurons (Fig 1A, Movie S1). To confirm this and to investigate Golgi dynamics during peripheral neuron regeneration, dissociated cells were fixed at 2, 16, 24, 48 hours and 7 days post-injury and labelled for β-III-tubulin to label neuron specific microtubules and the cis Golgi marker GM130 (Fig 1B, B’). At 2 hours post-injury, the Golgi was fragmented and dispersed within the soma. Over the course of regeneration, the Golgi then compacted in the soma, most notably in the first 24 hours. Golgi compaction then stabilised between 24 hours and 7 days (number of disconnected components in the GM130 channel quantified in Fig 1B’’’; 3 animals, 44 cells at 2h, 50 cells at 16h, 36 cells at 24h, 49 cells at 48h, 43 cells at 7 days). To characterise how these conformational changes in the Golgi related to the stages of regeneration we determined the number of cells at 2, 16, 24, 48 hours and 7 days post-injury exhibiting various hallmarks of regeneration (Fig S1; 3 animals). At 2 hours the majority of cells either possessed clear retraction bulbs (38/76 cells), an indication of acute injury, or had no extensions (37/76) while a single cell displayed an asymmetric reduction in microtubule labelling that we term the ‘microtubule penannular’ (Fig S1A). At 16 hours there was a reduction in the number of cells with retraction bulbs (24/90) or with no extensions (27/90), while the number of cells with a microtubule penannular increased (36/90). A small number of these (3/90) now also possessed multiple extensions. At 24 hours there was a further reduction in cells with retraction bulbs (16/102) or no extensions (17/102), and a corresponding increase in cells with a microtubule penannular (60/102) or multiple extensions (5/102). Furthermore, a small number of these cells now displayed a dominant axon (4/102). This trend continued at 48 hours, with no cells displaying a retraction bulb (0/80), fewer cells with no extensions (2/80) or a microtubule penannular (6/80). Conversely, there was an increase in cells with multiple extensions (43/80) or a dominant axon (29/80). By 7 days most cells either displayed multiple extensions (32/62) or a dominant axon (30/62) (Fig S1B). We then correlated these morphological hallmarks of regeneration with the state of Golgi compaction in these cells. This revealed that cells with clear retraction bulbs exhibited the greatest number of disconnected Golgi components, closely followed by cells with no extensions and cells at the microtubule penannular stage. In contrast, cells with multiple extensions or with a dominant axon displayed a sharp decrease in the number of disconnected Golgi components, demonstrating a correlation between Golgi compaction and initiation of axon regeneration (quantified in Fig S1C).

**Figure 1:**
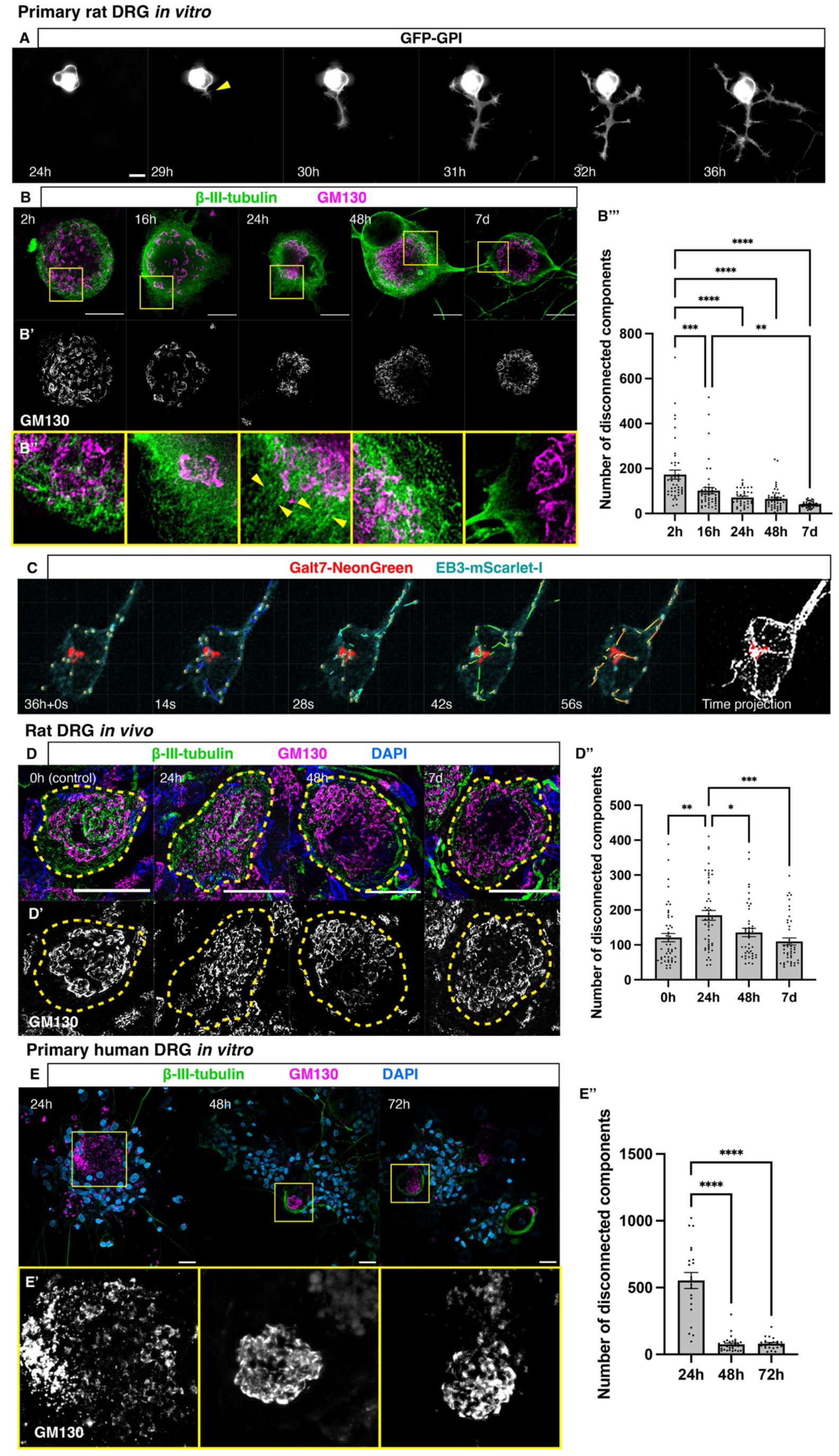
Golgi Compaction is associated with emergence of microtubules from the Golgi and initiation of axon regeneration. **A)** Time-lapse sequence of dissociated DRG neuron expressing GFP-GPI initiating axon regeneration at 29 hours post-injury (yellow arrowhead). **B)** Immunostaining to detect β-III-tubulin and GM130 from 2 hours up to 7 days post-injury, imaged using super-resolution STED microscopy. **B’)** GM130 shown in grayscale, highlighting the changes in Golgi morphology from 2 hours to 7 days post-injury. Yellow boxes in (B) outline zoomed regions in **B’’**), where linear tracks of microtubules appear to emerge from the Golgi at 24 hours (yellow arrowheads). **B’’’)** Quantification of mean number of disconnected components in the GM130 channel: 173 ± 20.26 at 2 hours, 101.5 ± 13.94 at 16 hours, 70.94 ± 6.12 at 24 hours, 65.02 ± 6.72 at 48 hours and 40.19 ± 1.97 at 7 days p=0.0003 2 vs 16 hours; p<0.0001 2 hours vs 24 hours, 48 hours and 7 days; p=0.003 16 hours vs 7 days. **C)** Time lapse sequence of dissociated DRG neuron at 36 hours post-injury transfected with Galt7-NeonGreen and EB3-mScarlet-I. Tracks display the trajectory of EB3 comets emerging from Golgi. Time-projected image displays microtubule tracks over the entire duration of the movie. **D)** Immunolabelling of tissue sections from L4/5 rat DRGs following sciatic nerve transection and subsequent recovery for 0 hours, 24 hours, 48 hours and 7 days for β-III-tubulin and GM130. Yellow dashed lines outline individual cells. **D’)** GM130 shown in grayscale demonstrating fragmentation at 24 hours post-injury followed by compaction to baseline (control) from 48 hours to 7 days. **D’’)** Quantification of mean number of disconnected components in GM130 channel: 121 ± 11.95 at 0 hours, 184.6 ± 3.97 at 24 hours, 135.7 ± 11.94 at 48 hours and 109.7 ± 9.95 at 7 days; p=0.0012 0 hours vs 24 hours, p=0.0251 24 hours vs 48 hours, p=0.0001 24 hours vs 7 days. **E)** Primary human DRG in vitro labelled for β-III-tubulin and GM130. Yellow boxes outline zoomed regions in **E’)** GM130 shown in grayscale, highlighting the changes in Golgi morphology from 24 hours to 72 hours post-injury. **E’’)** Quantification of mean number of disconnected components in GM130 channel: 553.1 ± 60.07 at 24 hours, 75 ± 8.93 at 48 hours and 80.33 ± 9.65 at 72 hours; p=<0.0001 24 hours vs 48 hours, p=<0.0001 24 hours vs 72 hours. Scale bars: 20mm. All graphs displayed as mean ± SEM,*p≤0.05, **p≤0.01, ***p≤0.001, ****p≤0.0001; ordinary one-way ANOVA and Tukey’s post hoc test used for statistical analyses.

We further observed using super-resolution stimulated emission depletion (STED) microscopy (Fig 1B’’) that microtubules exhibited a disorganised appearance at 2- and 16-hours post-injury. Conversely, at 24 hours, when axon regeneration initiates, linear tracks of microtubules were visualised that characteristically appeared to emerge from the Golgi, suggesting that the compacted Golgi may be acting as a platform for microtubule nucleation at this timepoint. It appeared, however, that this was transient, as somatic microtubules appeared to return to their disorganised state at 48 hours and 7 days post-injury. To determine the origin of nascent microtubules in regenerating peripheral neurons, we monitored microtubule nucleation patterns by labelling microtubule plus-ends with EB3-mScarlet-I and the Golgi with Galt7-NeonGreen in dissociated DRG neurons *in vitro*. Live imaging at 36 hours post-injury, when expression of transfected constructs was established, revealed microtubule comets emerging from the Golgi and entering the axonal compartment (Fig 1C, Movie S2; 13 cells, 4 animals). Furthermore, time projections facilitated visualisation of microtubule tracks (Fig 1C, far right panel), confirming their origin from the Golgi. Taken together, these results strongly suggest that Golgi compaction corresponds with Golgi-mediated microtubule nucleation during initiation of peripheral neuron regeneration *in vitro*.

To confirm the physiological relevance of these findings, we performed *in vivo* unilateral sciatic nerve transections on adult rats with post-injury survival periods of 0, 24, 48 hours and 7 days. At each timepoint, animals underwent terminal anaesthesia and immediate isolation and fixation of L4/L5 DRG, which are the major contributors to the sciatic nerve. Fixed DRG were sectioned and labelled with β-III-tubulin and GM130 (Fig 1D, D’; 3 animals, 48 cells at 0h, 50 cells at 24h, 44 cells at 48 h, 46 cells at 7 days). This revealed that the Golgi *in vivo* exhibited similar responses to injury as dissociated neurons *in vitro*. In the control condition where DRG were immediately extracted from the animal and fixed (0 hours), the Golgi remained condensed within the soma. In contrast, by 24 hours post-injury a dispersal pattern was evident similar to that observed at 2 hours *in vitro*. By 48 hours, the Golgi had condensed and remained as such until 7 days. These observations were confirmed by disconnected component analysis (Fig 1D’’). Therefore, DRG neuron Golgi *in vivo* exhibit a similar response to injury as dissociated neurons *in vitro*, but the *in vivo* response proceeds at a slower pace.

We then explored the clinical relevance of these findings by fixing and labelling human donor-derived DRG neurons *in vitro* with β-III-tubulin and GM130 at 24, 48 and 72h post-injury (Fig 1E, E’; 3 human donors, 21 cells at 24h, 34 cells at 48h, 21 cells at 72h). As human derived DRG neurons are a limited resource, we based these timepoints on our previous observation in rats *in vivo*. This revealed similar Golgi dynamics to those observed in rat DRGs *in vitro and in vivo*. The Golgi was fragmented and dispersed within the soma at 24 hours post-injury followed by compaction at 48 hours post-injury and maintenance of Golgi compaction at 72 hours post-injury (quantified in Fig 1E’’). These findings indicate that the pattern of injury-induced Golgi fragmentation followed by Golgi compaction is conserved between rats and humans.

### Microtubule nucleation factors AKAP9 and γ-tubulin associate with the Golgi during initiation of axon regeneration

Golgi mediated microtubule nucleation is facilitated by recruitment of AKAP9 and γ-tubulin to the cis-Golgi membrane in several experimental systems, including cultured cells and Drosophila neurons (16-18). To investigate if similar mechanisms operate in the context of adult vertebrate peripheral neuron regeneration, we labelled dissociated rat DRG neurons for the cis-Golgi protein GM130, AKAP9 and γ-tubulin. This revealed AKAP9 puncta associated with the Golgi at all time-points from 2 hours to 7 days post-injury, suggesting recruitment of AKAP9 to the Golgi at all stages (Fig 2A; 115 cells at 2h, 128 cells at 16h, 88 cells at 24h; 46 cells at 48h, 51 cells at 7 days; 3 animals). In contrast, labelling for γ-tubulin and GM130 revealed that although γ-tubulin was expressed at all time points, there was no apparent association with the Golgi at 2 hours, 16 hours and 7 days (Fig 2B; 62 cells at 2h, 117 cells at 16h, 151 cells at 7 days; 3 animals). Strikingly, we observed clear association at 24 and 48 hours, which respectively correspond with Golgi compaction (Fig 1A; 85 cells at 24h, 51 cells at 48h; 3 animals) and emergence of microtubules from the Golgi (Fig 1C), suggesting that recruitment of γ-tubulin at these specific timepoints may facilitate Golgi-mediated microtubule nucleation. To further investigate this, we performed proximity ligation assays (PLA) to confirm interactions between GM130, AKAP9 and γ-tubulin from 16 hours to 7 days post-injury. This confirmed recruitment of AKAP9 to the Golgi at all stages, but we observed fewer and more intense PLA puncta from 24 hours post-injury, suggesting that aggregation of Golgi-associated AKAP9 following Golgi compaction corresponds with initiation of Golgi-mediated microtubule nucleation (Fig 2A’, A’’; 27 cells at 16h, 40 cells at 24h, 29 cells at 48h, 37 cells at 7 days; 2 animals). Consistent with this, we observed minimal recruitment of γ-tubulin to the Golgi at 16 hours, robust recruitment at 24 hours post-injury followed by a sharp decline in recruitment at 48 hours which persisted until 7 days post-injury (Fig 2B’, B’’; 32 cells at 16h, 25 cells at 24h, 27 cells at 48 h, 28 cells at 7 days; 2 animals). The specificity of these experiments was confirmed by performing PLA on cells fixed at 24 hours post-injury that had not been labelled for GM130 and AKAP9 or γ-tubulin, which resulted in no detectable PLA puncta (Fig S2; 24 cells; 2 animals).

**Figure 2:**
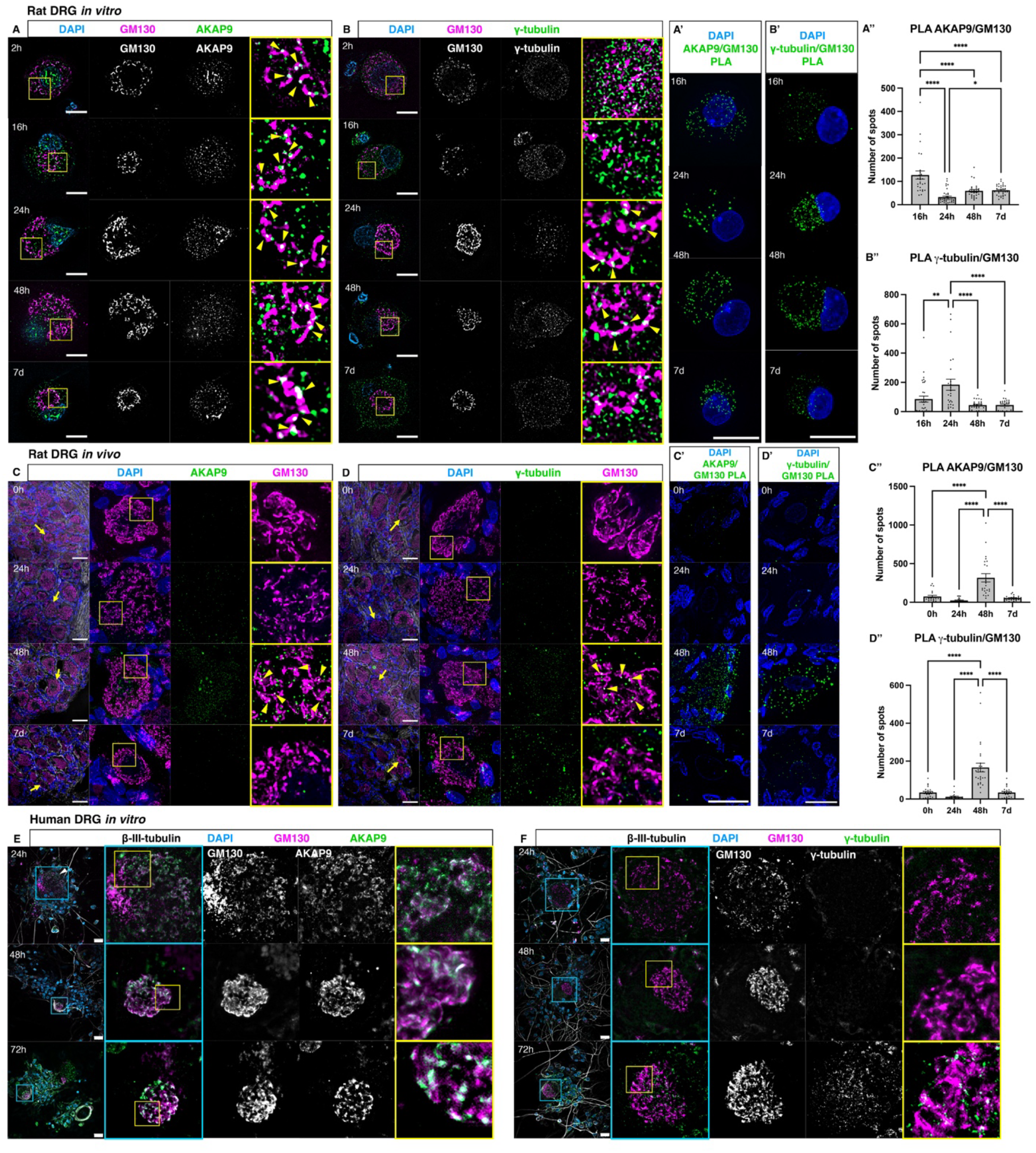
Sequential recruitment of AKAP9 and γ-tubulin to the compacting Golgi. **A)** Immunolabelling of dissociated DRG neurons 2 hours to 7 days post-injury for GM130 and AKAP9. Yellow arrowheads: AKAP9 puncta decorating Golgi. Yellow boxes outline zoomed in regions. **A’)** PLA to determine interactions between GM130 and AKAP9 at 16 hours to 7 days. **A’’)** Quantification of PLA puncta: 127.3 ± 17.52 at 16 hours; 32.68 ± 4.29 at 24 hours; 59.55 ± 5.62 at 48 hours and 61.76 ± 3.48 at 7 days. p<0.0001 16 vs 24 hours, 48 hours and 7 days; p=0.0381 24 hours vs 7 days. **B)** Immunolabelling of dissociated DRG neurons 2 hours to 7 days post-injury for GM130 and γ-tubulin. Yellow arrowheads: γ-tubulin puncta decorating Golgi. Yellow boxes outline zoomed in regions. **B’)** PLA to determine interactions between GM130 and γ-tubulin at 16 hours to 7. **B’’)** Quantification of PLA puncta: 85.81 ± 20.64 at 16 hours, 184.2 ± 37.75 at 24 hours, 44.48 ± 5.97 at 48 hours and 44.96 ± 7.07 at 7 days. p=0.0069 16 hours vs 24 hours; p<0.0001 24 hours vs 48 hours and 7 days. **C)** Immunolabelling of L4/5 rat DRGs tissue sections following sciatic nerve transection and recovery for 0/24/48 hours and 7 days for GM130 and AKAP9. Yellow arrowheads: AKAP9 puncta decorating Golgi. Zoomed in regions identified by yellow arrows/boxes. **C’)** PLA to determine interactions between GM130 and AKAP9 at 0 hours to 7 days post-injury. **C’’)** Quantification of PLA puncta: 74.9 ± 16.13 at 0 hours, 22.67 ± 6.23 at 24 hours, 316.8 ± 53.37 at 48 hours and 53.91 ± 6.13 at 7 days; p<0.0001 48 hours vs 0 hours, 24 hours and 7 days. **D)** Immunolabelling of tissue sections for GM130 and γ-tubulin. Yellow arrowheads: γ-tubulin puncta decorating Golgi. **D’)** PLA to determine interactions between GM130 and γ-tubulin at 0 hours to 7 days post-injury. **D’’)** Quantification of PLA puncta: 34.32 ± 6.16 at 0 hours, 1.6 ± 3.51 at 24 hours, 166.3 ± 23.09 at 48 hours and 34.32 ± 6.16 at 7 days; p<0.0001 48 hours vs 0 hours, 24 hours and 7 days. **E)** Dissociated primary human DRG fixed at 24/48/72 hours post-injury, labelled for GM130 and AKAP9. Zoomed in regions enclosed in blue/yellow boxes. **F)** Dissociated primary human DRG fixed at 24/48/72 hours post-injury, labelled for GM130 and γ-tubulin. Zoomed in regions enclosed in blue/yellow boxes. Scale bars: 20mm. All graphs display mean± SEM.*p≤0.05, **p≤0.01, ****p≤0.0001; ordinary one-way ANOVA and Tukey’s post hoc test used for statistical analyses.

In the *in vivo* context of rat sciatic nerve transections, we observed clear AKAP9 and γ-tubulin association with the Golgi at 48 hours post-injury, coinciding with Golgi compaction (Fig 1D), but not at 0 hours, 24 hours or 7 days post-injury (Fig 2C, D; Fig 2C, D; 48 cells at 0h, 50 cells at 24h, 44 cells at 48h, 46 cells at 7 days post-injury; 3 animals). These associations were confirmed by PLA, which revealed robust recruitment of both AKAP9 (Fig 2C’, C’’; 20 cells at 0h, 21 cells at 24h, 23 cells at 48h, 23 cells at 7 days post-injury; 2 animals) and γ-tubulin (Fig 2D’. D’’; 22 cells at 0h, 20 cells at 24h, 28 cells at 48h, 22 cells at 7 days post-injury; 2 animals) to the Golgi at 48 hours post-injury, but not at 0 hours, 24 hours or 7 days post-injury.

To determine if similar dynamic recruitment of AKAP9 and γ-tubulin takes place in the context of human peripheral neuron regeneration, we fixed and labelled human donor-derived DRG neurons *in vitro* with GM130, AKAP9 and γ-tubulin at 24-, 48- and 72-hours post-injury. Similar to our findings *in vitro* in rat, we observed clear associations between the Golgi and AKAP9 at all time-points (Fig 2E; 46 cells at 24h, 44 cells at 48h, 39 cells at 72h; 3 donors). At 24- and 48-hours post-injury, no associations between the Golgi and γ-tubulin were evident (Fig 2F; 47 cells at 24h, 60 cells at 48h; 3 donors). However, at 72 hours post-injury, we observed a clear association between the Golgi and γ-tubulin puncta (Fig 2F; 48 cells; 3 donors). Taken together, these observations strongly suggest that Golgi compaction following peripheral neuron injury is accompanied by recruitment of AKAP9 to the compacted Golgi, which in turn facilitates recruitment of the key microtubule nucleation factor γ-tubulin. Notably, this pattern of stepwise recruitment is conserved between rat and human species but proceeds at a slower pace in the context of human PNI.

### An intact Golgi is required to facilitate microtubule nucleation and drive axon regeneration

To substantiate the role of the Golgi as an MTOC during peripheral neuron regeneration, we applied Brefeldin A (BFA) which inhibits protein transport from endoplasmic reticulum (ER) to Golgi and so disassembles the Golgi (19). In doing so, we assessed whether an intact, compacted Golgi is a prerequisite for microtubule nucleation and subsequent axon regeneration.

BFA was applied to dissociated rat DRG cells at 23 hours post-injury, when the Golgi is compacted (Fig 1B’), followed by fixation 1 hour later. Labelling for GM130 and AKAP9 or γ-tubulin revealed that application of BFA induced disruption of the Golgi compared to cells grown in medium containing DMSO (Fig 3A, B, quantified in Fig 3C; 30 cells BFA treated, 30 cells DMSO treated; 2 animals). Furthermore, induced disruption of the Golgi also resulted in loss of AKAP9 and γ-tubulin association with GM130 (Fig 3A, B, far right panels; 37 cells γ-tubulin labelled, 39 cells AKAP9 labelled; 2 animals). These observations were confirmed by PLA for interactions between GM130 and AKAP9 or γ-tubulin (Fig 3A’, B’, quantified in Fig 3A’’, B’’; AKAP9/GM130: 32 DMSO cells, 20 BFA treated cells; 2 animals. γ-tubulin/GM130: 34 DMSO cells, 24 BFA treated cells; 2 animals), indicating that an intact, compacted Golgi is required for recruitment of these key microtubule nucleating factors. BFA was then applied to cells expressing EB3-mScarlet-I to label microtubule plus-ends and Galt7-NeonGreen to label the Golgi at 36 hours post-injury followed by timelapse imaging. This revealed rapid disruption of the Golgi within 10 minutes and an associated decrease in microtubule polymerisation events, evidenced by a gradual loss of EB3 comets at 10 minutes and 30 minutes following application of BFA (Fig 3D, Movie S3; 3 animals, 8 cells). Longer-term timelapse imaging of cells expressing GFP-GPI immediately after addition of BFA over 15 hours revealed that disruption of the Golgi resulted in errors in axon regeneration in the majority of cells imaged (15/16 cells, 2 animals) (Fig 3E, Movie S4). Of these, 10/16 were unable to initiate axon regeneration (Movie S4) and 5/16 possessed axons at the point of BFA application, all of which subsequently retracted towards the cell body. To confirm that these effects were caused by reduced microtubule nucleation, and to discount the effect of impaired Golgi-mediated trafficking, we applied the γ-tubulin functional inhibitor Gatastatin G2 (20). This resulted in disrupted axon regeneration, where all treated cells were unable to initiate axonal outgrowth (11/11 cells, 2 animals) (Fig 3E, Movie S5). This contrasted with cells imaged in medium containing DMSO, the majority of which were able to undergo axon regeneration (15/23 cells, 2 animals) (Fig 3E, Movie S6). To confirm these findings in cells that were not subjected to our live-imaging regime, we fixed BFA treated cells at 40 hours post-injury and labelled for GM130 and β-III-tubulin (Fig 3F). This again revealed disruption of the Golgi compared to DMSO treated cells, indicating that the effects of BFA application persist at 16 hours post-application (quantified in Fig 3F’; BFA treated: 30 cells; DMSO treated: 30 cells; 2 animals). Furthermore, most BFA treated cells displayed errors in axon regeneration (18/21 cells, 2 animals). Of these, 6/21 exhibited no regeneration, 12/21 had retraction bulbs present in neurites, 1/21 had multiple extensions and 2/21 had an obvious axon (Fig 3F’’). Similarly, Gatastatin G2 treated cells also exhibited errors in axon regeneration (39/40 cells, 2 animals) (Fig 3F), but the Golgi remained compacted in these cells (quantified in Fig 3F’). Of these, 33/40 cells exhibited no regeneration, 6/40 had retraction bulbs present, 1/40 had multiple extensions and 0/40 had an obvious axon (Fig 3F’’). This contrasted with cells grown in medium containing DMSO, in which 7/42 exhibited no regeneration, 0/42 had retraction bulbs, 23/42 multiple extensions and 12/42 an obvious axon (Fig 3F’’). These results indicate that an intact, compacted Golgi is a prerequisite for recruitment of AKAP9 and γ-tubulin, leading to Golgi-mediated microtubule nucleation that drives axon regeneration following injury.

**Figure 3:**
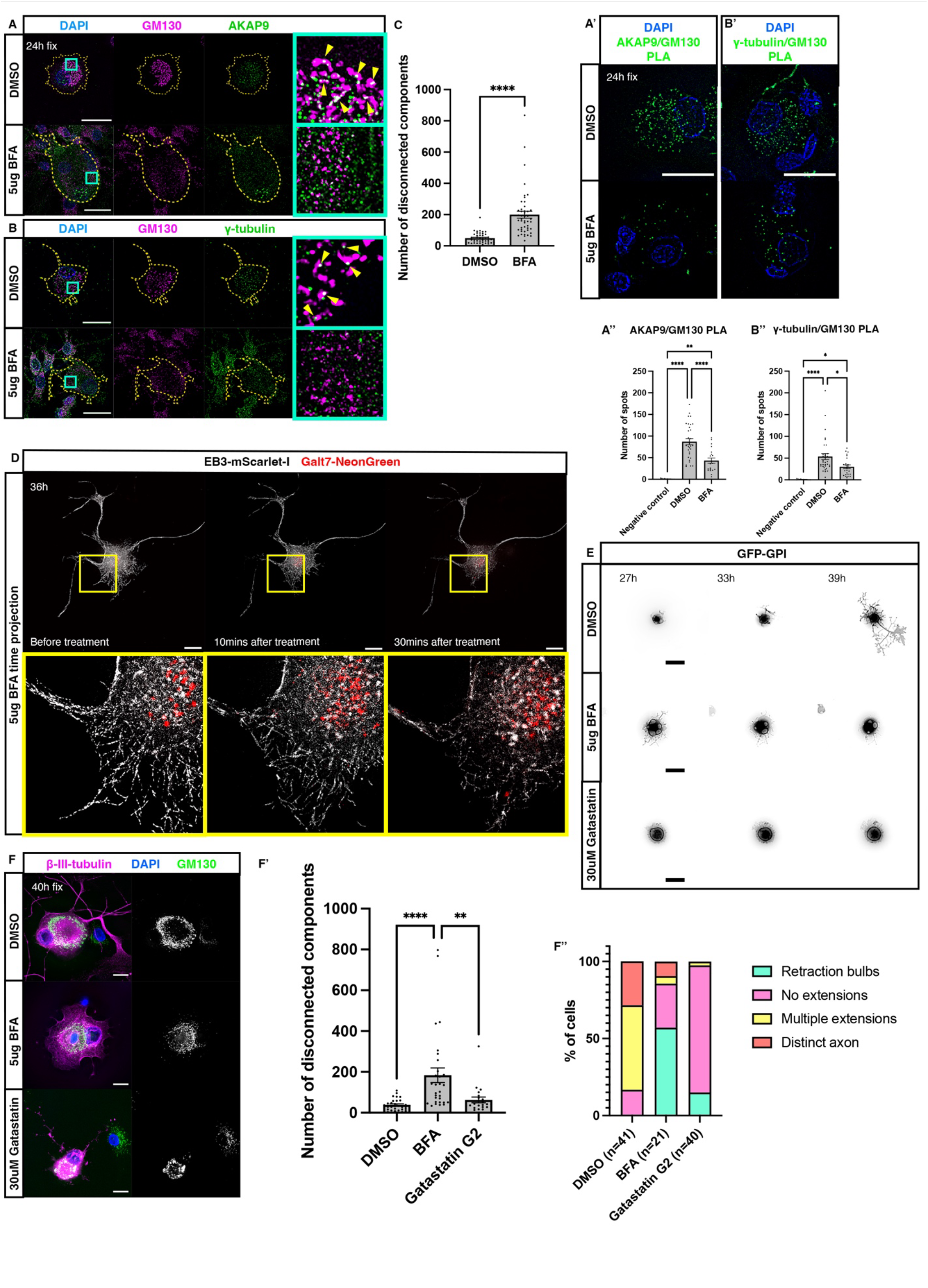
Golgi compaction is required for initiation of axon regeneration. **A)** Immunostaining to detect GM130 and AKAP9 in dissociated DRG neurons cultured in medium containing DMSO (top panels) or BFA (bottom panels). Yellow arrowheads: AKAP9 puncta decorating Golgi. Blue boxes indicate zoomed in regions shown in right-hand panel. **B)** Immunostaining to detect GM130 and γ-tubulin in dissociated DRG neurons cultured in medium containing DMSO (top panels) or BFA (bottom panels). Yellow arrowheads: γ-tubulin puncta decorating Golgi. Blue boxes indicate zoomed in regions shown in right-hand panel. **A’)** PLA to determine interactions between GM130 and AKAP9 in dissociated DRG neurons treated with DMSO or BFA. **A’’)** Quantification of PLA puncta: 87.06 ± 7.09 DMSO and 43.25 ± 6.1 BFA treated cells. **B’)** PLA to determine interactions between GM130 and γ-tubulin in dissociated DRG neurons treated with DMSO or BFA. **B’’)** Quantification of PLA puncta: 53.56 ± 6.87 DMSO (34 cells) and 30.17 ± 4.29 BFA treated cells. **C)** Quantification of mean number of disconnected components in GM130 channel: 49.56 ± 5.13 in DMSO treated cells and 199.3 ± 21.92 in BFA treated cells; p=<0.0001. **D)** Time projections of 1-minute timelapse sequence of cell expressing EB3-mScarlet-I and Galt7-NeonGreen following addition of BFA. Yellow boxes indicate zoomed in regions shown in bottom panels. **E)** Timelapse sequences of cells expressing GFP-GPI imaged in medium containing DMSO (top panels), BFA (middle panels) and Gatastatin G2 (bottom panels). **F)** Immunolabelling of cells cultured in media containing DMSO (top panel), BFA (middle panel) or Gatastatin G2 (bottom panel) and fixed at 40 hours to detect β-III-tubulin and GM130. **F’)** Quantification of mean number of disconnected components in GM130 channel: 38.87 ± 5.1 in DMSO treated cells, 184 ± 35.9 in BFA treated cells and 63.23 ± 4.1 in Gatastatin G2 treated cells; p<0.0001 DMSO vs BFA treated cells; p=0.0027 Gatastatin G2 treated cells vs BFA treated cells. **F’’)** Quantification of the percentage of cells at observed stages of regeneration for each treatment. Scale bars: 20mm. All graphs displayed as mean ± SEM. *p≤0.05, **p≤0.01, ***p≤0.001, ****p≤0.0001; ordinary one-way ANOVA and Tukey’s post hoc test used for statistical analyses.

### Golgi associated AKAP9 is required for recruitment of γ-tubulin to drive initiation of axon regeneration

To determine if Golgi associated AKAP9 is required for recruitment of γ-tubulin and axon regeneration, cells were transfected with a construct expressing an AKAP9 fragment that competitively displaces AKAP9 from the Golgi (AKAP9-dis) (21). At 24 hours post-injury *in vitro*, AKAP9-dis displaced endogenous AKAP9 away from the Golgi compared to non-transfected cells (Fig 4A; 7 transfected cells, 7 non-transfected controls; 2 animals). This was accompanied by a striking displacement of γ-tubulin away from the Golgi compared to non-transfected cells, resulting in a scattered cellular distribution of γ-tubulin puncta (Fig 4B; 9 transfected cells, 4 non-transfected cells; 2 animals). These observations were confirmed by proximity ligation assays, which demonstrated loss of interactions between GM130 and AKAP9 (Fig 4A’; 56 cells; 2 animals) or γ-tubulin (Fig 4B’; 27 cells; 2 animals) in cells expressing AKAP9-dis compared to non-transfected cells (88 cells AKAP9/GM130 and 85 cells γ-tubulin/GM130; 2 animals) indicating that Golgi associated AKAP9 is required for recruitment of γ-tubulin. We then investigated if failure to recruit γ-tubulin to the compacted Golgi leads to errors in initiation of axon regeneration by performing CRISPR mediated knockouts of AKAP9. Cells were transfected with an AKAP9 targeting CRISPR knock-out construct expressing cytoplasmic blue fluorescent protein (BFP) (px458-AKAP9-BFP), fixed at 48 hours post-injury and labelled for AKAP9 and GM130 (Fig S3). This revealed a clear reduction of AKAP9 expression in knockout cells compared to non-transfected cells (9 transfected cells, 24 non-transfected controls; 1 animal). We then subjected cells transfected with px458-AKAP9-BFP and GFP-GPI to live imaging starting at 24-48 hours post injury. While the majority of cells not transfected with px458-AKAP9-BFP initiated axon regeneration (34/40 cells, 5 animals), cells lacking AKAP9 were unable to initiate axon regeneration up to 64 hours post injury (34/36 cells, 5 animals) (Fig 4C, Movies S7, S8 and Fig S4, Movies S9, S10), suggesting that recruitment of AKAP9 followed by γ-tubulin to the compacted Golgi is required for initiation of axon regeneration. Taken together, these results demonstrate that recruitment of AKAP9 to the compacted Golgi is followed by recruitment of γ-tubulin, which then drives Golgi mediated microtubule nucleation to facilitate initiation of axon regeneration.

**Figure 4:**
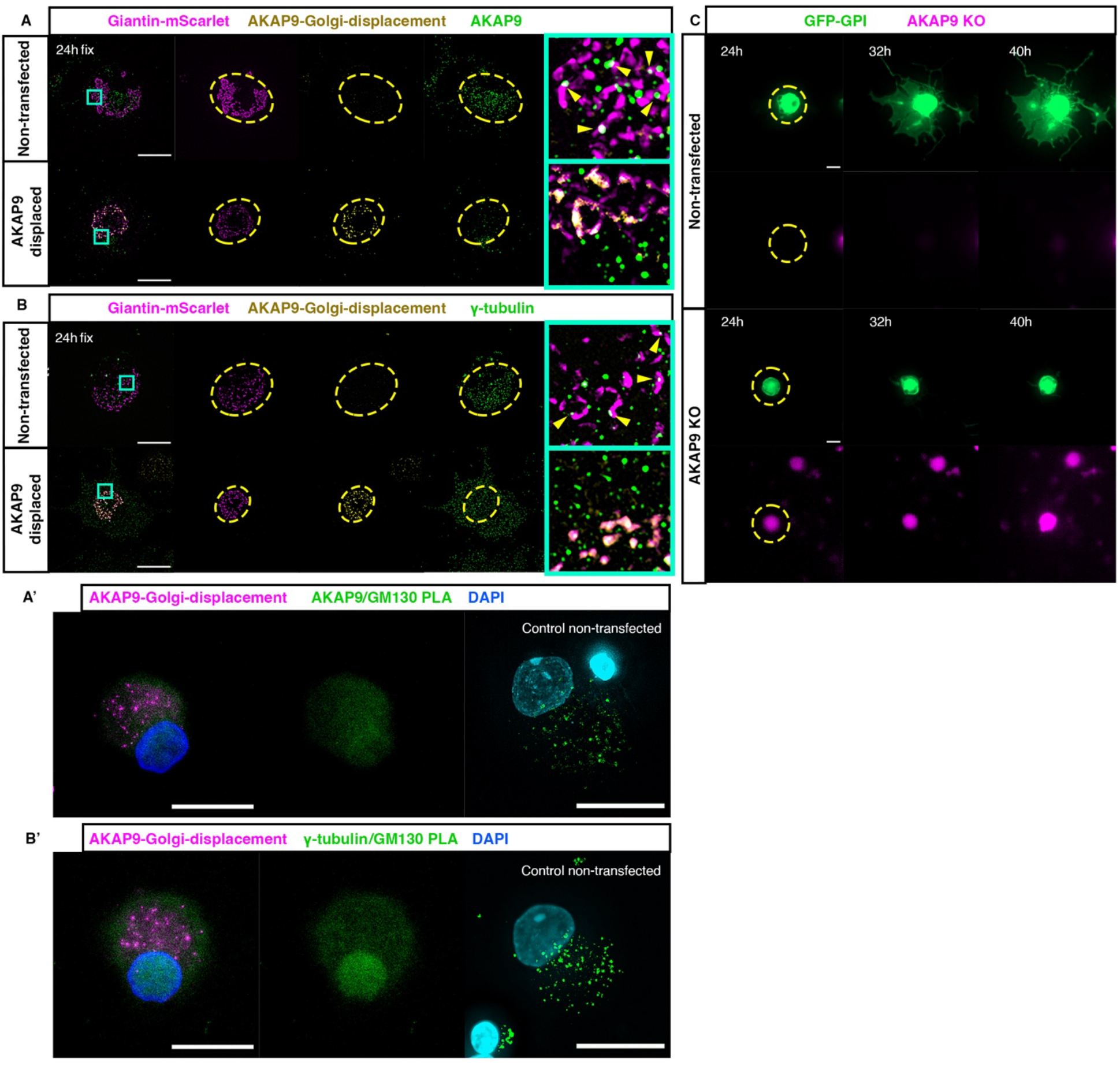
AKAP9 is required for recruitment of γ-tubulin to the Golgi and subsequent axon regeneration. **A)** Cells fixed at 24 hours expressing cis/medial Golgi marker Giantin-mScarlet and AKAP9-dis and immunostained for endogenous AKAP9. Top panel shows control cells not expressing AKAP9-dis and bottom panel cells expressing AKAP9-dis. Yellow dashes outline the area in the cell occupied by the Golgi and cyan boxes outline zoomed in regions. Yellow arrowheads: AKAP9 puncta decorating Golgi. **A’)** PLA to determine interactions between GM130 and AKAP9 at 24 hours, with non-transfected control cells in the rightmost panel. **B)** Cells fixed at 24 hours expressing Giantin-mScarlet, AKAP9-dis and immunostained for endogenous γ-tubulin. Top panel shows control cells not expressing AKAP9-dis and bottom panel cells expressing AKAP9-dis. Yellow dashes outline the area in the cell occupied by the Golgi and cyan boxes outline zoomed regions. Yellow arrowheads: γ-tubulin puncta decorating Golgi. **B’)** PLA to determine interactions between GM130 and γ-tubulin at 24 hours, with control cells in the rightmost panel. **C)** Timelapse sequences of cells expressing GFP-GPI not expressing AKAP9 knockout construct (top panels) and expressing AKAP9 knockout construct (magenta, bottom panels).

## Discussion

We demonstrate that peripheral nerve injury (PNI) triggers dramatic fragmentation of the somatic Golgi. This is followed by rapid compaction of the Golgi and recruitment of the scaffolding protein AKAP9, which then recruits the key microtubule nucleating factor γ-tubulin during a discrete period. This regulated sequence of events ultimately results in microtubule nucleation from the compacted somatic Golgi which drives initiation of axon regeneration (Fig 5). Importantly, this mechanism is conserved between human and rat species, confirming the clinical relevance of these findings in PNI. Consistent with previous reports indicating that peripheral neuron regeneration proceeds at a slower pace in larger animals (22,23), we report distinct differences in the timing of Golgi dynamics in rat and human neurons *in vitro*, with a 24-hour delay in Golgi fragmentation in the human context. Furthermore, compared to cells *in vitro*, Golgi fragmentation is delayed by 24 hours in the context of *in vivo* sciatic nerve transections, which are performed distal to the DRG cell bodies (4 centimetres). This delay is consistent with proximal injuries inducing a more rapid neuronal response and the reported rate of retrograde axon transport (24-26), suggesting that our observed Golgi response is initiated by the second wave of injury signalling via retrograde transport of injury-responsive transcription factors rather than the rapid retrograde calcium wave which determines neuron survival (27).

**Figure 5:**
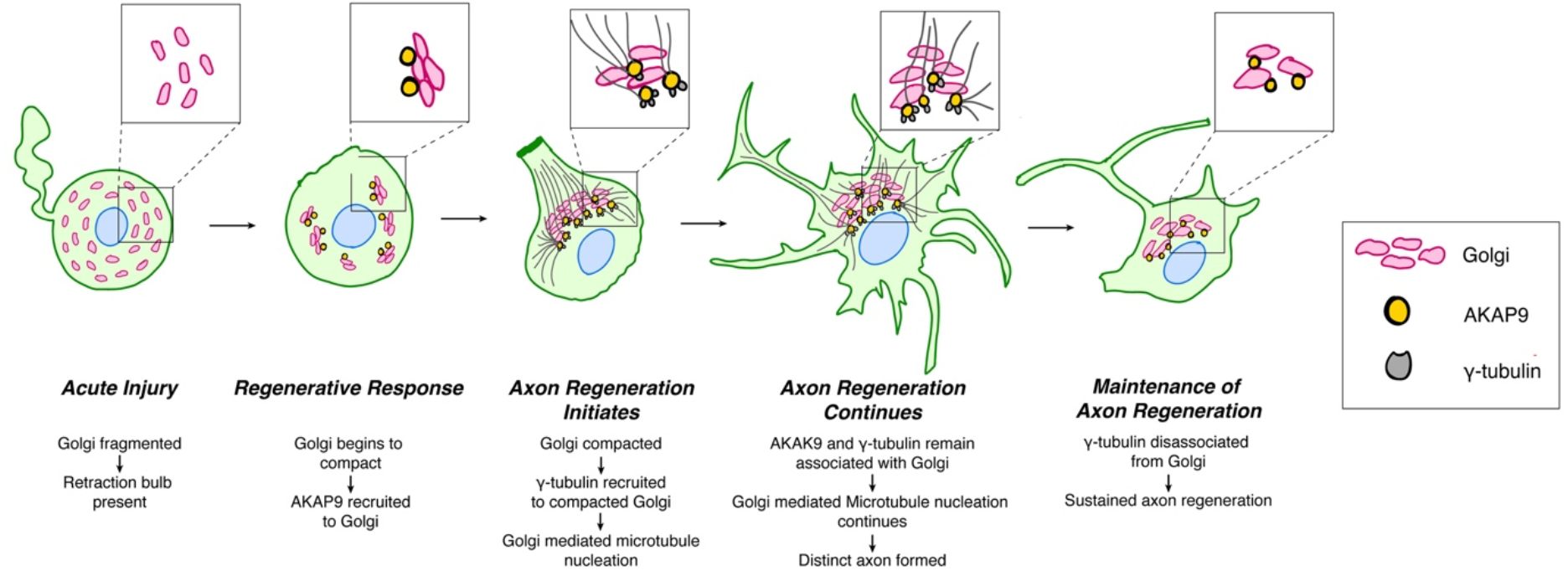
Schematic outlining the sequence of events driving initiation of peripheral neuron regeneration. Peripheral neuron injury results in fragmentation and subsequent compaction of the Golgi. Golgi compaction coincides with recruitment of AKAP9, which then facilitates recruitment of γ-tubulin and leads to Golgi mediated microtubule nucleation to drive initiation of axon regeneration. Following initiation of axon regeneration, γ-tubulin disassociates from the Golgi as axon regeneration continues.

We further report that AKAP9 recruitment to the somatic Golgi precedes that of γ-tubulin, supporting evidence that AKAP9 enhances recruitment of γ-tubulin to the Golgi (16,18, 28). Notably, γ-tubulin recruitment to the Golgi is transient *in vitro and in vivo*, and coincides with initiation of axon regeneration, following which γ-tubulin is again excluded from the somatic Golgi. This finding raises the possibility that, in the acute state of injury, the somatic Golgi mediated response initiates regeneration of an axon, but once this process is underway, alternative MTOC(s) coordinate maintenance of axon regeneration, perhaps through further acentrosomal nucleation sites within the regenerating axon itself. These findings therefore determine a critical temporal window during which injured peripheral neurons are competent to initiate regeneration. Novel pharmacological interventions delivered through retrograde signalling alongside timely surgical repair could potentiate more neurons at earlier timepoints to switch to a compacted Golgi and microtubule nucleating phenotype. Priming the innate neuronal regenerative response in this way would maximise the potential for clinically meaningful recovery of nerve function.

The phenomenon of Golgi fragmentation followed by rapid compaction in injured peripheral neurons is of particular interest as various neurodegenerative disorders, including Alzheimer’s disease and Parkinson’s disease, are hallmarked by fragmentation of the Golgi prior to presentation of clinical symptoms (29,30). Intriguingly, neuronal repair mechanisms are impeded in these disorders, and Golgi fragmentation in this context is followed by cell death, implicating Golgi fragmentation as an indicator of cellular insult, introducing the possibility that subsequent compaction of the fragmented Golgi may be a key first step towards initiation of repair mechanisms. In agreement with this concept, we demonstrate that induced disruption of an already compacted Golgi results in reduced AKAP9 and γ-tubulin recruitment, compromised Golgi-mediated microtubule nucleation and failed initiation of axon regeneration, further indicating that maintenance of Golgi architecture is required to sustain the initial steps of regeneration. Although the mechanisms promoting Golgi compaction in the context of peripheral neuron regeneration remain unclear, a key difference in comparison to diseased neurons of the central nervous system is the proximity of satellite glial cells that promote survival of injured peripheral neurons through expression of neurotropic factors, including NGF, BDNF and NT-3 (31). Consistent with this, we have observed clusters of non-neuronal likely glial cells associated with regenerating peripheral neurons at the microtubule penannular stage becoming the site of axon initiation in vitro (Fig S1). This suggests that signalling from these cells may promote survival of injured peripheral neurons by facilitating Golgi compaction and determining axon initiation.

Intriguingly, maintenance of Golgi structure itself relies on the cellular actin and microtubule network (32-34), suggesting that compaction of the Golgi during the regenerative response is interlinked with the pre-existing cytoskeletal pool. This raises the possibility that polarised cytoskeletal remodelling in response to extracellular signalling cues may facilitate compaction of the fragmented Golgi and trigger the regenerative response. DRG neurons display a differential response to injury, where neurons with smaller cell bodies are more likely to undergo cell death (35,36). It is therefore of particular importance to study this variable response to injury to determine if Golgi fragmentation is a reliable indicator of cell death, and if induction of Golgi compaction through potentiation of extracellular signalling cues can improve the regenerative response in populations of peripheral neurons.

In addition to its role as a platform for microtubule nucleation, it is abundantly clear that the Golgi performs several additional roles that are likely to contribute to the regenerative response. As a key mediator of cellular protein trafficking, Golgi positioning controls polarised membrane transport and therefore maintains cells in a polarised state (37). It is therefore conceivable that Golgi compaction during peripheral neuron regeneration may also facilitate the polarised expansion of the cell membrane which is characteristic of axon extension. Furthermore, polarised trafficking from the compacted Golgi is also likely to deliver key cellular receptors to the regenerating axonal compartment, facilitating a polarised response to extracellular signalling cues.

Overall, this work identifies fragmentation followed by compaction of the somatic Golgi as an initiator of axon regeneration in injured peripheral neurons. These findings identify the Golgi as a potential target for therapeutic interventions aiming to potentiate the scale and rate of peripheral neuron regeneration and may inform our understanding of the mechanisms that accelerate progression of neurodevelopmental disorders.

## Materials and Methods

### Ethics statement

Adult rats have been utilised in this study, having undertaken appropriate Home Office training. All experimental procedures adhere to the principles of replacement, reduction and refinement for use of animals in biological research and in accordance with the United Kingdom Animals (Scientific Procedures) Act 1986 under project licence PP9645155.

### Rat dorsal root ganglia dissection and dissociation

Adult Sprague Dawley rats (between 8-12 weeks of age) were sacrificed by carbon dioxide asphyxiation followed by cervical dislocation. Spinal columns were dissected away from the animal through a dorsal longitudinal incision and subsequent steps undertaken in a laminar flow hood. Muscle and connective tissue were dissected away from the vertebral column before midline longitudinal cuts were made down the length of the vertebral column in a cranio-caudal direction on both the ventral and dorsal surfaces, such that the spinal cord and vertebral foramen were exposed. The spinal cord was gently dissected away using micro forceps under microscopic guidance. DRG were then extracted from each vertebral foramen by firm but controlled traction. Extracted DRGs were placed immediately in ice cold F12 medium supplemented with 1% penicillin/streptomycin in a 70 mm petri dish. Each animal yielded approximately 40 DRG. Following extraction DRG nerve roots were micro-surgically divided to leave only the body of the DRG remaining. Connective tissue and excess blood were also removed. DRGs were then dissociated to single cells for our *in vitro* model (38).

All dissociation steps were carried out in a biological safety cabinet (class II. Explanted and cleaned DRGs were placed in a 30 mm petri dish containing 1.8 mL warm F12 medium. 200μl 1.25% wt/v type IV collagenase (Gibco) was then added and DRG incubated at 37°C for 60 minutes. The medium was removed and replaced with fresh F12 medium containing collagenase. DRG were incubated for a further 45 minutes at 37°C. Collagenase solution was removed and DRG washed three times with 2mL warm F12. 1.8mL F12 was then added along with 200μl 2.5% wt/v trypsin (Gibco) and DRG incubated for 30 minutes at 37°C. Trypsin was then removed and 1.5 mL F12 with 500μl foetal bovine serum (FBS) was added to stop trypsinisation. DRG were again washed three times with 2 mL F12 and after the final wash a further 2 mL F12 was added and DRG transferred using a Pasteur pipette to a 15 mL falcon tube. DRG were triturated by gently pipetting up and down using a Pasteur pipette 8 to 10 times. Remaining undissociated ganglia/debris were allowed to settle and supernatant containing dissociated cell suspension transferred to a fresh 15 mL falcon. A further 2 mL F12 was added to undissociated DRG and the process of trituration and supernatant transfer repeated until all DRG were dissociated. The resultant dissociated cell suspension was passed through a sterile 70 μm mesh to remove debris then centrifuged at 900 rpm for 5 minutes at 37°C. To achieve a neuron-rich culture a 15% bovine serum albumin (BSA) solution was prepared by combining 500 μl BSA with 500 μl F12. This solution was pipetted down the side of a clean 15 mL falcon such that a ‘track’ was formed over which the cell solution would be applied. After centrifugation, the supernatant was removed, and cells resuspended in 500 μl F12. The cell suspension was pipetted into the BSA containing falcon along the previously formed ‘track’ and centrifuged at 1200 rpm for 10 minutes at 37°C. Cells were then either resuspended in nucleofector solution, in the case of transfection (see below), or in an appropriate volume of F12 supplemented with N2 for immunofluorescence staining. Cells were plated onto either 13 mm 1.5 glass coverslips in 4-well plates or FluoroDish 35 mm dishes ready coated with poly-D-lysine (PDL) (World Precision Instruments). Glass coverslips were coated with 0.1 ug/ul PDL for 20 minutes and washed once with F12. Glass coverslips and/or FluoroDishes were then coated with laminin, prepared 1:100 in F12 (to reach a final concentration of 1-2 μg/cm^2^). Plates/dishes were incubated for 2 to 24 hours at 37°C. After coating, the laminin solution was removed, and glass washed once with F12 prior to cell plating.

### Dissociated rat DRG electroporation

Dissociated DRG were resuspended in 100 μl Lonza nucleofector solution (brought to room temperature 30 minutes before use). For optimal transfection 1 million cells were required, which equates to the DRG from one whole rat. A total of 1-2 μg (1 μg per construct) DNA plasmid solution was then added to the cell suspension and gently mixed by pipetting up and down. The suspension was then transferred to a sterile Lonza nucleofection cuvette, ensuring that the suspension contained no air bubbles and was at the bottom of the cuvette. Programme G-013 was selected on the Nucleofector device and the cuvette inserted into the carrier before performing electroporation. Following electroporation, 500 μl of warmed supplemented media was added to the cuvette and transferred carefully using the supplied Lonza plastic Pasteur pipette to a sterile Eppendorf tube. The resultant suspension was then incubated at 37°C for 10 minutes to aid cell recovery from electroporation. During the recovery time warmed supplemented media was applied to prepared coverslips/FluoroDishes (to result in 150 μl per 13 mm coverslip and 600 μl per FluoroDish) and these dishes incubated at 37°C until plating. Without repeated aspirations, the cell suspension was then divided between the prepared plates/ FluoroDishes (4 times the volume of suspension was added to FluoroDishes when compared to 13 mm coverslips). The cells were incubated at 37°C for 2 hours, after which media containing residual nucleofector solution was removed and fresh supplemented media added (F12 with 1:100 N2; 600 μl per well in four well plates and 2 mL per FluoroDish). Media was subsequently changed three times per week.

### Plasmids

The following plasmids were used: pCAGGS-GFP-GPI, Galt7-NeonGreen, pCAG-mScarlet-Giantin (a gift from Frank Bradke, Addgene plasmid # 196872^21^), EB3-mScarlet-I (a gift from Dorus Gadella, Addgene plasmid # 98826) (39), pRD04b-AKAP9g1, pRD04b-AKAP9g2, pRD04b-AKAP9g3 (AKAP9 CRISPR knock out constructs designed and produced in collaboration with The University of Manchester genome editing core facility), pBactin-AcGFP-128-425-Akap9 (a gift from Frank Bradke, Addgene plasmid # 196871^21^) at a concentration between 500 ng -1 μg per construct, not exceeding 2 μg per transfection.

### *In vivo* rat sciatic nerve injury

Adult male Lewis rats between 8-12 weeks of age were used for *in vivo* study due to the lower incidence of autotomy following nerve transection. All animals were acclimatised for at least 1 week prior to surgery. Animals were anaesthetised using isofluorane inhalation anaesthesia. Induction was carried out in an induction chamber with 2 L/minute oxygen flow and 5% isofluorane until animals lost their righting reflex and breathing slowed. Maintenance of anaesthesia was achieved using a nose cone and 2-2.5% isofluorane. Throughout surgery depth of anaesthesia was monitored regularly by assessing breathing rate/depth and pedal withdrawal reflex. 20 minutes prior to the initiation of surgery, all animals received subcutaneous buprenorphine analgesia (0.3 mg/mL stock) at a dose of 0.01 mg/kg and the surgical area was minimally shaved. Surgery was carried out using an aseptic technique and animals placed on a 37°C heat pad to minimise heat/fluid loss. The sciatic nerve was exposed through a dorsal gluteal approach and transection performed at the level of the proximal femur. Animals were allowed to recover for: 0 hours, 24 hours, 48 hours and 7 days before undergoing terminal anaesthesia by overdose of isofluorane followed by cervical dislocation. L4 and L5 DRG were then extracted and immediately fixed in 4% PFA for 4 hours at room temperature.

### Primary human dorsal root ganglia

Human DRG were obtained from AnaBios (USA). Cadaveric DRG from consenting donors with no comorbidities were extracted, dissociated, cultured and fixed by the suppliers according to their optimised protocols before being shipped to our lab. Cells were cultured on glass bottom 96-well plates on which immunofluorescence labelling was performed.

### Fixation and immunofluorescence labelling of dissociated DRG cells

Dissociated DRGs were fixed at the following time points: 2 hours, 16 hours, 24 hours, 48 hours and 7 days (rodent cells) and 24 hours, 48 hours and 72 hours (human cells) post-injury. For each time-point, experimental triplicates were used. Cells were fixed with 4% paraformaldehyde (PFA) for 20 minutes at room temperature. Following fixation, cells were washed once with phosphate buffered saline (PBS) for 5 minutes. Prior to staining, PBS was removed and cells permeabilised using 0.2% Triton X100 for 30 minutes at room temperature, followed by a PBS wash. Blocking was then carried out using normal donkey serum (Abcam) diluted 5:100 in antibody diluent (prepared by adding 1 g BSA, 1 g sodium azide and 300 μl Triton X-100 to 1 L PBS). Following blocking cells were then incubated with primary antibodies (table 1) for either 2 hours at room temperature or 16 hours at 4°C. Cells were then washed in PBS once for 5 minutes. PBS was removed and secondary antibodies (table 1) diluted 1:500 in antibody diluent were applied and plates incubated at room temperature for 1 hour in the dark. Following incubation, cells were washed with PBS once and counterstained for DAPI (1:1000) for 3 minutes followed by a final PBS wash. Coverslips were mounted onto glass Superfrost™ slides (Thermo Fisher Scientific) using either ProLong™ Diamond or ProLong Glass antifade mounting medium (Thermo Fisher Scientific) slides/dishes were stored at 4°C both prior to, and following, imaging.

**Table 1:**
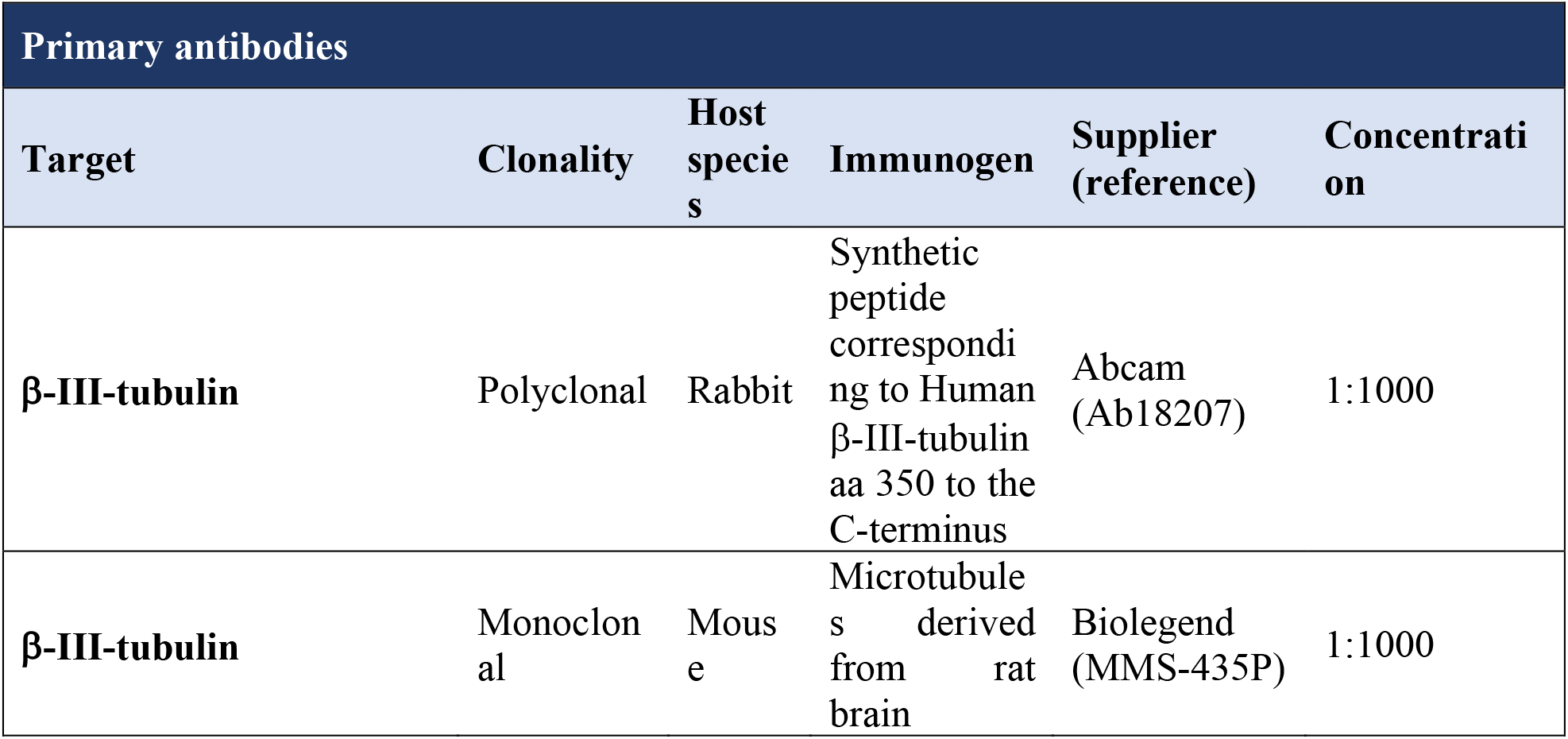
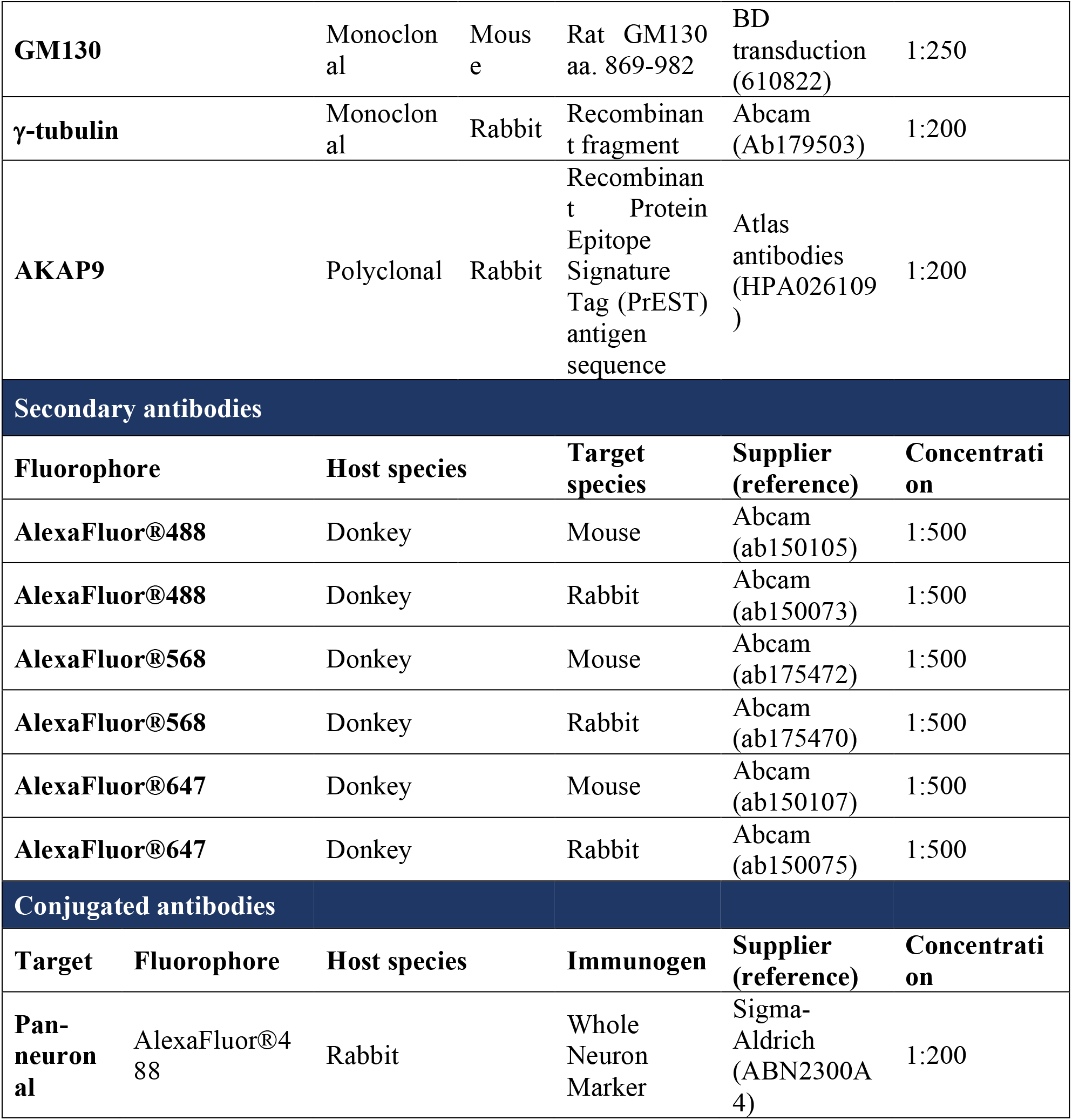
Antibodies used in this work including suppliers and concentrations used.

### Fixation and immunofluorescence labelling of DRG explants

L4 and L5 DRG from *in vivo* experiments were extracted and immediately fixed in 4% PFA for 4 hours at room temperature. DRG were washed with PBS and then dehydrated in 30% sucrose solution for 24 hours (or until DRG sunk). They were then embedded in OCT and snap frozen on dry-ice before being placed at -80°C prior to sectioning. Sectioning was carried out on a Leica CM3050s cryostat and 20um sections obtained. Immunofluorescence staining was carried out as for *in vitro* cells.

## Table of antibodies

### Proximity ligation assays

Duolink Proximity Ligation Assay (PLA) kits were used according to manufacturer’s instructions. Fixed DRG on 13 mm glass coverslips were used at the following time-points: 16 hours, 24 hours, 48 hours and 7 days. Samples were first permeabilised using 0.2% Triton X100 for 30-minutes at room temperature. Blocking was then carried out using 40 μl Duolink blocking solution per coverslip. Coverslips were incubated in a humidity chamber at 37°C for 60 minutes. Blocking solution was then removed and replaced with 40μl primary antibodies (GM130 and AKAP9 or GM130 and γ-tubulin) diluted in Duolink antibody diluent. Samples were incubated in a humidity chamber for 2 hours at room temperature or 16 hours at 4°C. Following primary antibody incubation, the solution was removed, and coverslips washed twice with 1x Duolink wash buffer A at room temperature (5 minutes per wash). 40μl PLA PLUS and MINUS probes diluted 1:5 in Duolink antibody diluent were then added to each coverslip and samples incubated in a humidity chamber at 37°C for 60 minutes. Probes were then removed, and coverslips washed twice with wash buffer A. 1x Duolink ligation buffer was prepared by diluting the 5x buffer 1:5 with high purity water, then ligase added to the resultant solution at 1:40 immediately prior to addition to coverslips. Wash buffer was removed and 40 μl ligation solution was applied. Samples were incubated in a humidity chamber at 37°C for 30 minutes. Ligation solution was removed, and samples washed twice with wash buffer A. Amplification buffer was then prepared by diluting 5x amplification buffer 1:5 in high purity water and polymerase added immediately prior to use at 1:80 in amplification buffer. 40 μl of resultant solution was added to each coverslip and samples incubated for 100 minutes at 37°C. Amplification solution was removed, and samples washed twice with 1x wash buffer B at room temperature (10 minutes per wash). A final 1-minute wash was carried out using 0.01x wash buffer B. To prepare the samples for imaging, the remaining wash buffer was removed, and coverslips mounted onto glass slides using Duolink in situ mounting medium (containing DAPI) and edges sealed with clear nail varnish.

### Pharmacological treatments

*In vitro* DRG were treated at 23 hours post-injury and either underwent timelapse imaging or fixation after 1 hour of treatment for immunofluorescence staining with either 30 μM Gatastatin G2 (Funakoshi), 5 μg/mL BFA (Sigma-Aldrich), or control DMSO 1% diluted in F12 supplemented with 1:100 N2. In the case of EB3 imaging, cells were treated at 36 hours post-injury.

### Fixed cell and tissue imaging

Images were acquired using a Zeiss Cell Observer Z1 microscope system (Carl Zeiss) equipped with a Colibri7 light-emitting diode (LED) illumination system and 63× 1.4 numerical aperture (NA) objective and Flash4 v2 sCMOS camera (Hamamatsu). ZenPro 2.3 blue edition (Carl Zeiss) software was used for image acquisition and post-acquisition processing. Images were deconvolved using a constrained iterative algorithm, maximum 40 iterations and 0.1% quality threshold. Higher resolution was achieved for select cells using STED microscopy using a Leica Microsystems TCS SP8 STED system equipped with a 100× 1.4 NA oil immersion STED objective. Images were acquired using a 488 nm excitation laser and 592 nm depletion laser for the green channel and a 568 nm excitation laser and 660 nm depletion laser for the red channel. Z-sections were separated by 0.2 µm and images were scanned at 10 Hz using 2x line averaging. The resulting images were deconvolved using Huygens Professional (Scientific Volume Imaging).

### Time-lapse imaging

Time-lapse imaging was performed using a widefield Zeiss Axio-observer microscope system equipped with a Colibri7 LED illumination system (Carl Zeiss), Flash4 v2 sCMOS camera (Hamamatsu) and heated chamber with 5% carbon dioxide. For assessment of cell dynamics (GFP-GPI transfected cells), images were acquired every 10 to 20 minutes for 10 to 24 hours using a 40x 1.2 NA silicone immersion objective. For EB3 (EB3-mScarlet transfected cells) images were acquired as fast as possible for 1 to 3 minutes using a 63x 1.4 NA objective. Z plane intervals were 0.5um and minimal exposure times (20-50ms) were used in each channel. Time-lapse images were deconvolved in ZenPro using a fast iterative algorithm, 0.1% quality threshold and a maximum of 40 iterations.

### Image analysis

To perform analysis of Golgi disconnected components Imaris 9.2.1 software (Bitplane) was used. The surfaces function was used with automatic thresholding to reduce the risk of bias. This enabled 3-dimensional (3D) analysis of Golgi structure, including the number of discrete components, termed ‘number of disconnected components’.

Imaris was also used to quantify PLA puncta using the spot function. This enabled each distinct fluorescent signal to be picked up as a single spot. The number of spots and the volume of puncta occupying each cell were then quantified. The spot manual tracking function was also used to visualise EB3 comet trajectory.

### Statistical analysis

GraphPad Prism 9 (GraphPad Software, California, USA) was used to carry out statistical analysis. To quantify data for Golgi Apparatus disconnected components, as well as PLA spot analysis, a one-way analysis of variance (ANOVA) with Tukey’s multiple comparison test was used. P<0.05 was considered statistically significant.

## Acknowledgments

We thank V. Allan, M. Houslay, M. Lowe, K. Dorey, K. Storey and J. Wong for comments on the manuscript, P. March and S. Marsden from the University of Manchester Bioimaging Facility for technical support with microscopy and A. Adamson from the University of Manchester Genome Editing Unit for technical support with CRISPR.

## Author contribution

Conceptualisation and Supervision: RMD and AJR. Methodology: AEM, RMD, AJR. Investigation: AEM. Visualisation: AEM, RMD, AJR. Writing – original draft, review and editing: AEM, RMD, AJR.

## Data and materials availability

All data needed to evaluate the conclusions in the paper are present in the paper and/or the supplementary materials. Raw data files and all reagents may be requested from the authors.

