## Supplementary Figures for "Golgi compaction facilitates microtubule nucleation to drive adult vertebrate peripheral neuron regeneration"

Mortimer *et al.*

**This PDF file includes:**

Figs. S1 to S4
Movies S1 to S#

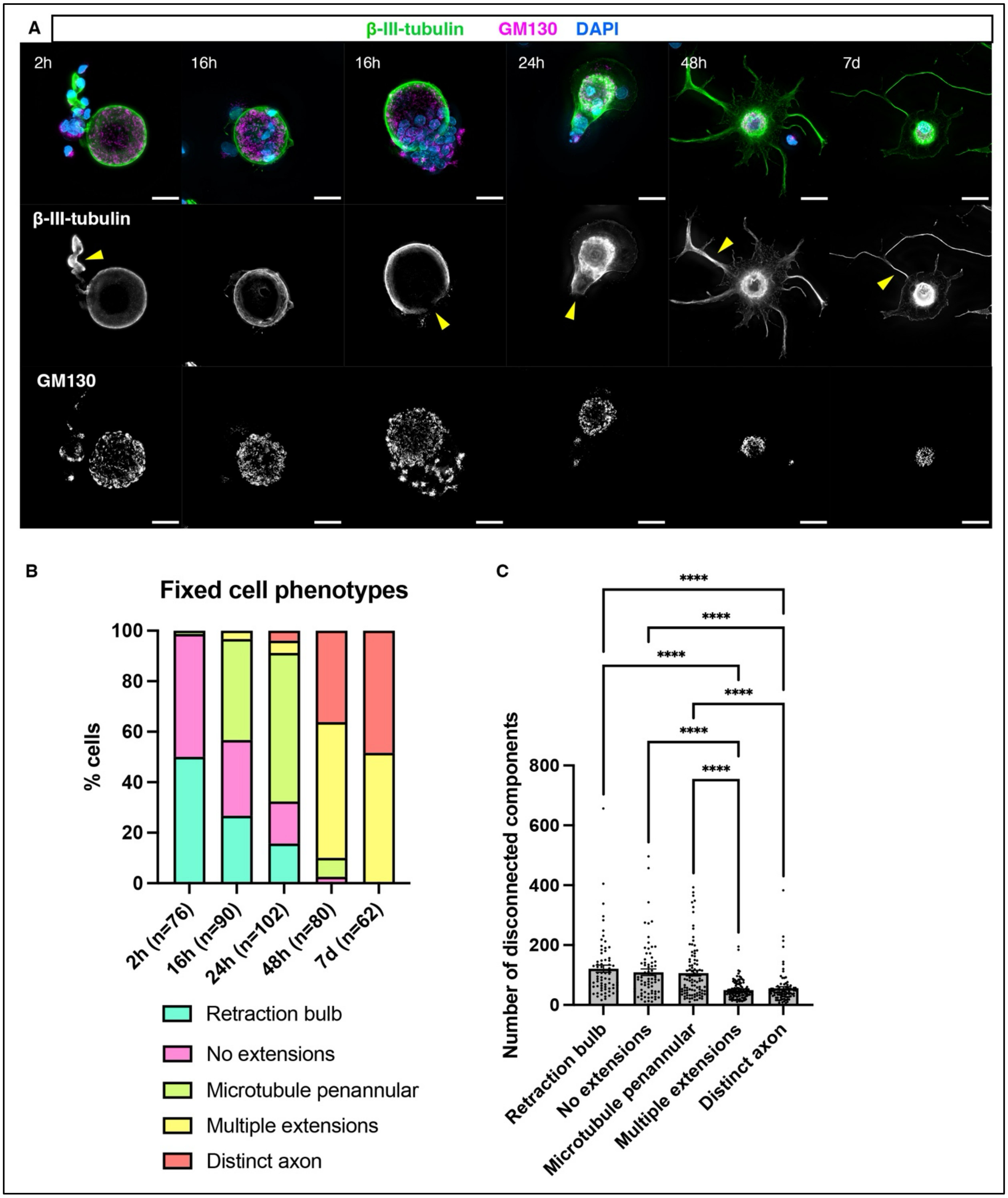

A
β-III-tubulin GM130 DAPI
2h
16h
16h
24h
48h
7d
β-III-tubulin
GM130
B
Fixed cell phenotypes
% cells
100
80
60
40
20
0
2h (n=76)
16h (n=90)
24h (n=102)
48h (n=80)
7d (n=62)
Retraction bulb
No extensions
Microtubule penannular
Multiple extensions
Distinct axon
C
****
Number of disconnected components
800
600
400
200
0
Retraction bulb
No extensions
Microtubule penannular
Multiple extensions
Distinct axon

**Fig. S1. Golgi Compaction is associated with initiation of axon regeneration.**

**A)** Immunostaining to detect neuron specific microtubules (β-III-tubulin) and GM130 from 2 hours up to 7 days post-injury. Yellow arrowheads: retraction bulb at 2 hours; the microtubule penannular at 16 hours; initiation of regeneration at 24 hours; multiple extensions at 48 hours; and a distinct axon at 7 days. **B)** Quantification of the percentage of cells at observed stages of regeneration at each time point. **C)** Quantification of mean number of disconnected components in GM130 channel according to cell phenotype: 120.8 ± 11.98 with retraction bulbs, 109.1 ± 1.08 with no extensions, 106.5 ± 9.26 with microtubule penannular, 49.36 ± 2.94 with multiple extensions and 55.34 ± 5.71 with a distinct axon. $p=<0.0001$ retraction bulb vs multiple extensions; $p=<0.0001$ retraction bulb vs distinct axon; $p=<0.0001$ no extensions vs multiple extensions; $p=<0.0001$ no extensions vs distinct axon; $p=<0.0001$ microtubule penannular vs multiple extensions; $p=<0.0001$ microtubule penannular vs distinct axon Scale bars: 20mm. **** $p \leq 0.0001$, ordinary one-way ANOVA and Tukey's post hoc test used for statistical analyses.

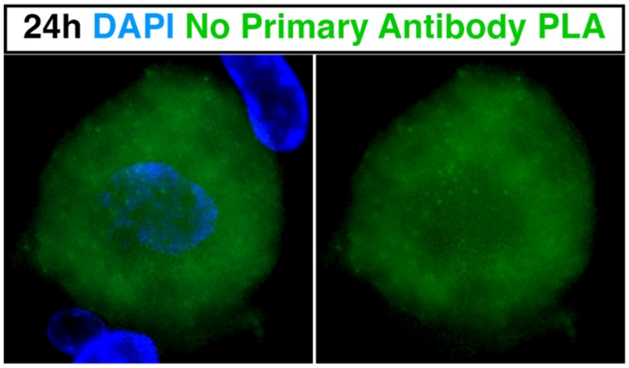


**Fig. S2. Negative control for proximity ligation assays. PLA carried out on cells at 24 hours post-injury in the absence of primary antibodies for GM130, AKAP9 or γ-tubulin.**

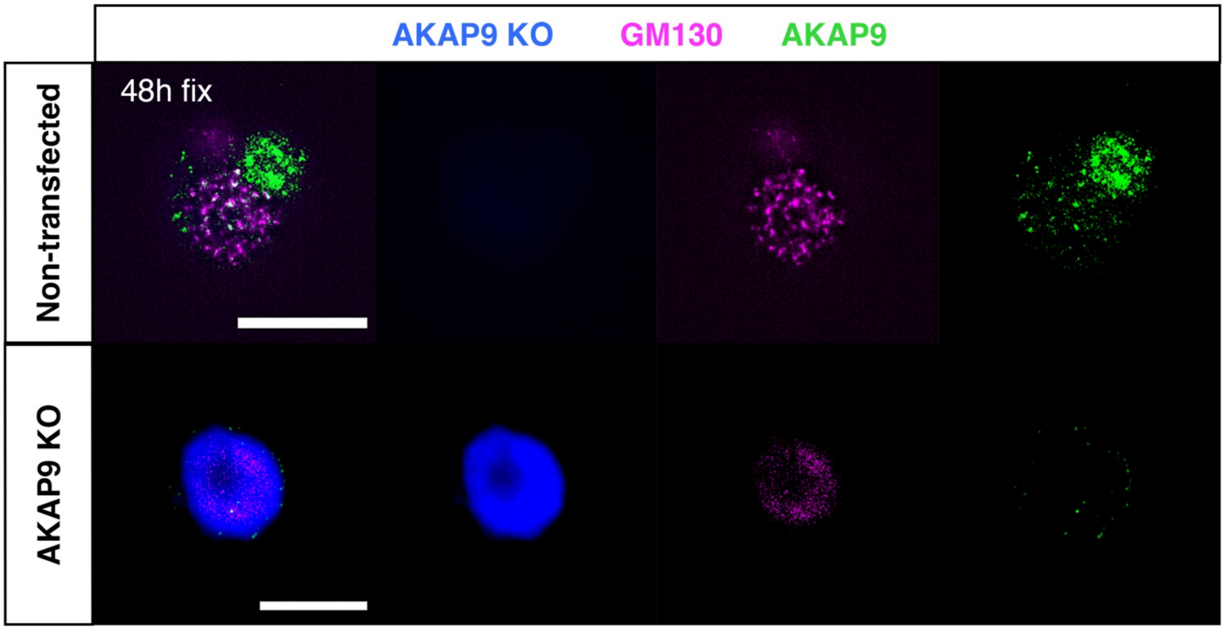


**Fig. S3. AKAP9 knock out reduces AKAP9 expression.** Control non-transfected cells (top panels) and AKAP9 KO cells (bottom panels) fixed at 48 hours post injury.

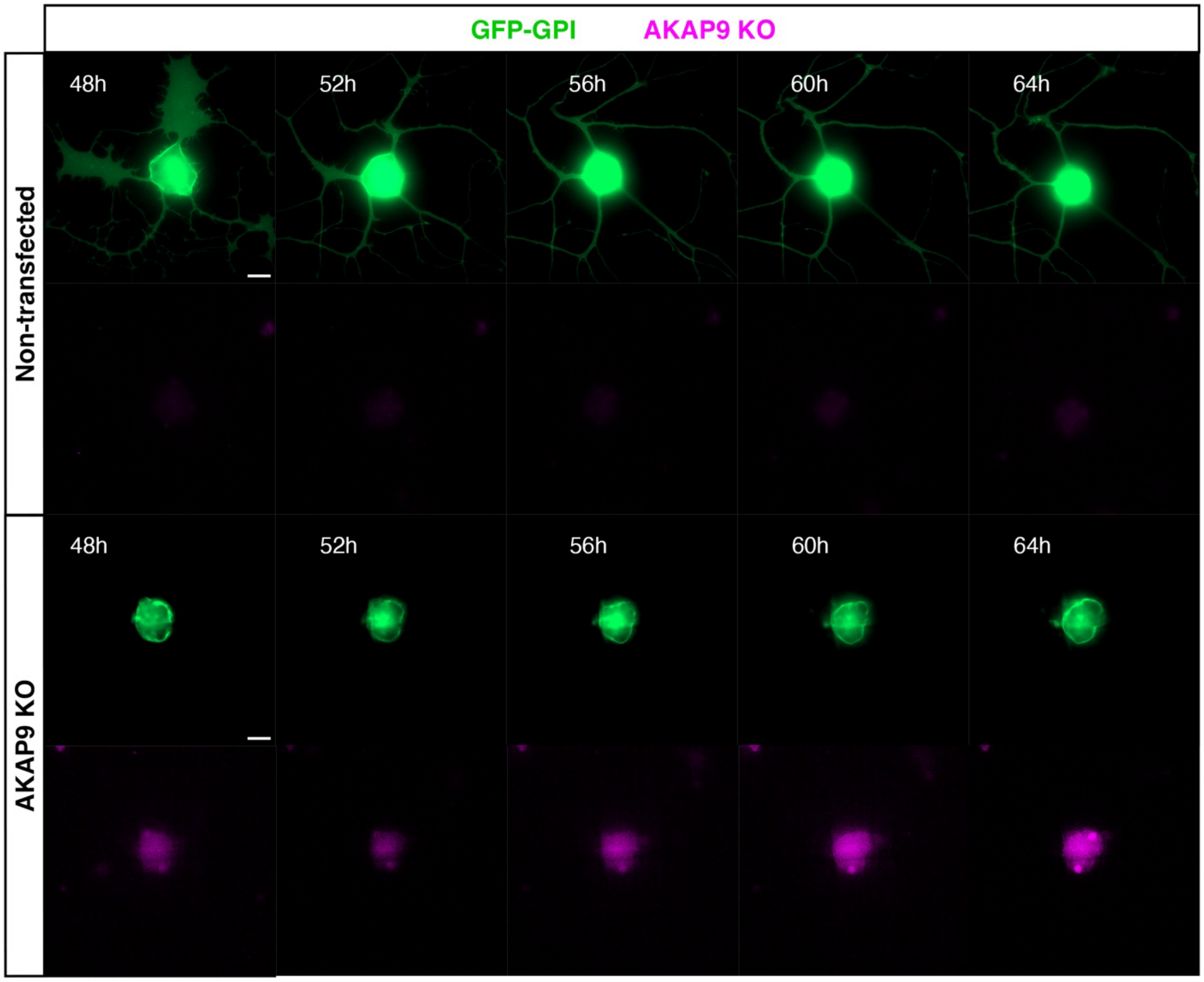


**Fig. S4. AKAP9 Knock out results in errors in axon regeneration from 48 to 64 hours following injury.** Timelapse images of cells expressing GFP-GPI to label the cell membrane (green) from 48 to 64 hours. Top panel shows a control cell extending an axon and bottom panel a cell expressing AKAP9 knock out construct (magenta).

**Movie S1.** Timelapse sequence of regenerating neuron expressing GFP-GPI (greyscale) at 24 – 40 hours post-injury.

**Movie S2**. Timelapse sequence of regenerating neuron expressing Galt7-NeonGreen (red) and EB3-mScarlet (cyan) imaged at 36 hours post injury for 2 minutes. EB3 + ends were tracked (yellow spheres and displacement tracks) and can be observed emerging from the Golgi.

**Movie S3.** Timelapse sequence of regenerating neuron expressing Galt7-NeonGreen (red) and EB3-mScarlet-I (greyscale) imaged at 36 hours post injury before a following application of BFA.

**Movie S4.** Timelapse sequence of regenerating neuron expressing GFP-GPI (green) imaged in medium containing BFA at 24 – 40 hours post-injury.

**Movie S5.** Timelapse sequence of regenerating neuron expressing GFP-GPI (green) imaged in medium containing Gatastatin G2 at 24 – 40 hours post-injury.

**Movie S6.** Timelapse sequence of regenerating neuron expressing GFP-GPI (green) imaged in medium containing DMSO at 24 – 40 hours post-injury.

**Movie S7.** Timelapse sequence of regenerating neuron expressing GFP-GPI (green) but not transfected with AKAP9 knockout construct at 24 – 40 hours post-injury

**Movie S8.** Timelapse sequence of regenerating neuron expressing GFP-GPI (green) and AKAP9 knockout construct (magenta) at 24 – 40 hours post-injury.

**Movie S9.** Timelapse sequence of regenerating neuron expressing GFP-GPI (green) but not transfected with AKAP9 knockout construct at 48 – 64 hours post-injury.

**Movie S10.** Timelapse sequence of regenerating neuron expressing GFP-GPI (green) and AKAP9 knockout construct (magenta) at 48 – 64 hours post-injury.
